# Characterizing rhythmic wheel-turning behavioral patterns in cockroaches *Rhyparobia maderae* using machine learning

**DOI:** 10.64898/2026.09.11.747745

**Authors:** J. Olschewski, A. Brückner-Foit, N. Filimonova, H. Zolmon, M. Stengl, E. Friedmann

## Abstract

Organisms must adapt to environmental changes occurring across multiple time scales, with endogenous multiscale clocks coordinating physiology and behavior with recurring environmental rhythms, including the dominant 24-hour cycle and faster ultradian rhythms. The Madeira cockroach (*Rhyparobia maderae*) provides a suitable model for investigating such multiscale temporal organization. Here, locomotor activity was recorded in running-wheel experiments under constant darkness. While the endogenous circadian clock produces a clearly visible 24-hour rhythm, it remains unknown whether locomotor behavior also exhibits temporal patterns at additional time scales. These temporal patterns cannot be found by classical frequency analysis, as they are veiled by higher harmonics of the circadian rhythm which are in the same frequency range. Unsupervised machine learning methods such as K-Means clustering, self-organizing maps and Gaussian mixture are used in search for fast ultradian rhythms possibly linked to circadian cycles in locomotor activity. Prior to applying these methods, data metrics are defined which characterize bouts of activity (called activity impulses) compared to periods of reduced activity. A stochastic pattern was found in these activity metrics which characterizes the time distance between activity impulses. Across all approaches, a consistent ultradian rhythm of approximately one hour was identified in the timing of the activity maxima. This rhythm was mainly detected during the subjective night, suggesting circadian control, and appears to consist of two components with periods of approximately 40 minutes and 1.5 hours. The method proposed in this paper is applied to two cockroach groups with different levels of activity, and is generalizable to diverse datasets occurring in the form of a time series with a dominant rhythm.

**Author Summary:** Detecting and predicting regular environmental changes is essential for the survival of organisms. Geophysical rhythms, such as the 24-hour light–dark cycle caused by the Earth’s rotation, have therefore driven the evolution of endogenous circadian clocks. Besides the daily light–dark cycle, organisms also respond to faster and slower environmental rhythms, such as recurring food availability or mating opportunities, signaled via patterned chemical stimuli. These rhythms are likely more flexible than the circadian rhythm, allowing organisms to adapt to local conditions. Because they are shorter, more frequent, and less regular, they are harder to detect and remain poorly understood despite their roles in processes such as hormone regulation and brain function. The ancient species of the Madeira cockroach possesses multiscale endogenous clocks that regulate physiology and behavior. Using a machine learning framework, we detected hidden faster rhythms beyond the circadian cycle and identified a consistent temporal pattern in the intervals between activity bursts, pointing to a potential ultradian rhythm that had not previously been described. This framework may be applicable beyond cockroaches, enabling the detection of similar rhythms in other species and the analysis of temporal patterns in diverse time series datasets.

## 1. Introduction

### 1.1 General framework

Geophysical rhythms as external Zeitgeber provided the selection pressure for the evolution of endogenous biological clocks in organisms on earth. Endogenous biological clocks at multiple time scales autonomously orchestrate the timing of physiology and behavior of uni- and multicellular organisms, of plants and animals alike. They provide the basis for self-organized physiological homeostasis and allow for prediction of regular environmental changes generated by environmental Zeitgebers. Thus, multiscale endogenous clocks increase biological fitness and survival rates over the course of evolution. Best known are circadian clocks that generate endogenous rhythms with a period of about 24 hours, orchestrating the daily rest-activity cycle and allowing for entrainment to the environmental light-dark cycle [1–3]. In this study we investigate the behavior of Madeira cockroaches, an established model system in chronobiology, in search for endogenous clocks at multiple time scales generating interlinked behavioral rhythms [4,5].

Our goal is to identify locomotor activity rhythms at different frequencies, to examine whether and how they are interlinked over largely different time scales from msec to hours, a so far unsolved puzzle. To analyze the dynamics of internally generated rhythmic locomotor activity across different experimental settings, we use unsupervised machine learning. This paper is concerned with the development of methodology, i.e. selecting machine learning methods suitable for finding behavioral patterns in seemingly irregular locomotor activity data. Control animals with clearly apparent circadian rhythms were selected to search for possibly interlinked ultradian rhythms in their locomotor activity.

*Rhyparobia maderae* were maintained in mass colonies at the University of Kassel under a 12:12 light/dark cycle at ∼100 lx, 25 °C, and 50% relative humidity according to previously described conditions [6]. Colonies were kept exposed to the light regime in translucent, beige plastic bins (60 × 40 × 30 cm) containing litter and cardboard egg cartons as shelter. The cockroaches were provided with dried dog food, organic potatoes, apples, carrots, and water *ad libitum*. Only male cockroaches were employed for all experiments because their locomotor activity rhythm is more stable compared to females and in order to minimize contamination by escaping cockroaches.

Free-running locomotor activity of individual male cockroaches was recorded in custom-built running wheels in constant darkness (DD) at 25 °C and 50% relative humidity. Animals were housed individually one per wheel, with 16–20 wheels kept at appropriate distances (to prevent synchronization) together in a light-tight recording box, provided with food (rodent chow, ssniff V2144, Soest Germany) and water *ad libitum*. Wheels were made of transparent plastic (wall thickness ∼1 mm) with an outer diameter of 7 cm and an inner ring of 3.5 cm, giving a running channel approximately 3.5 cm wide. The outer ring was perforated with evenly spaced holes (∼4 mm diameter, ∼1.8 mm apart) to prevent frass and debris from obstructing locomotion. Two magnets mounted 180° apart on the outer rim allowed each half-rotation to be registered by a Hall sensor. Activity was logged by custom Arduino-based systems, and the sensor events were binned into 1-min intervals on the device and written to an SD card as plain text (.txt). For the present paper, these data were summarized in an EXCEL-file with the first column giving the time in minutes and the second column giving the number of turns by minute summed over all wheels. Only cockroaches showing stable locomotor activity rhythms for at least one week were kept for continuous running-wheel assays over several weeks.

Actograms [7] (Fig 1A) can be employed to visualize the activity rhythm and its disturbances (Fig 1A). The circadian rhythm can be monitored and its time shift is well visible. However, there is no straightforward way to evaluate ultradian rhythms which are not recognizable at first sight with this representation of the data.

**Fig 1.**
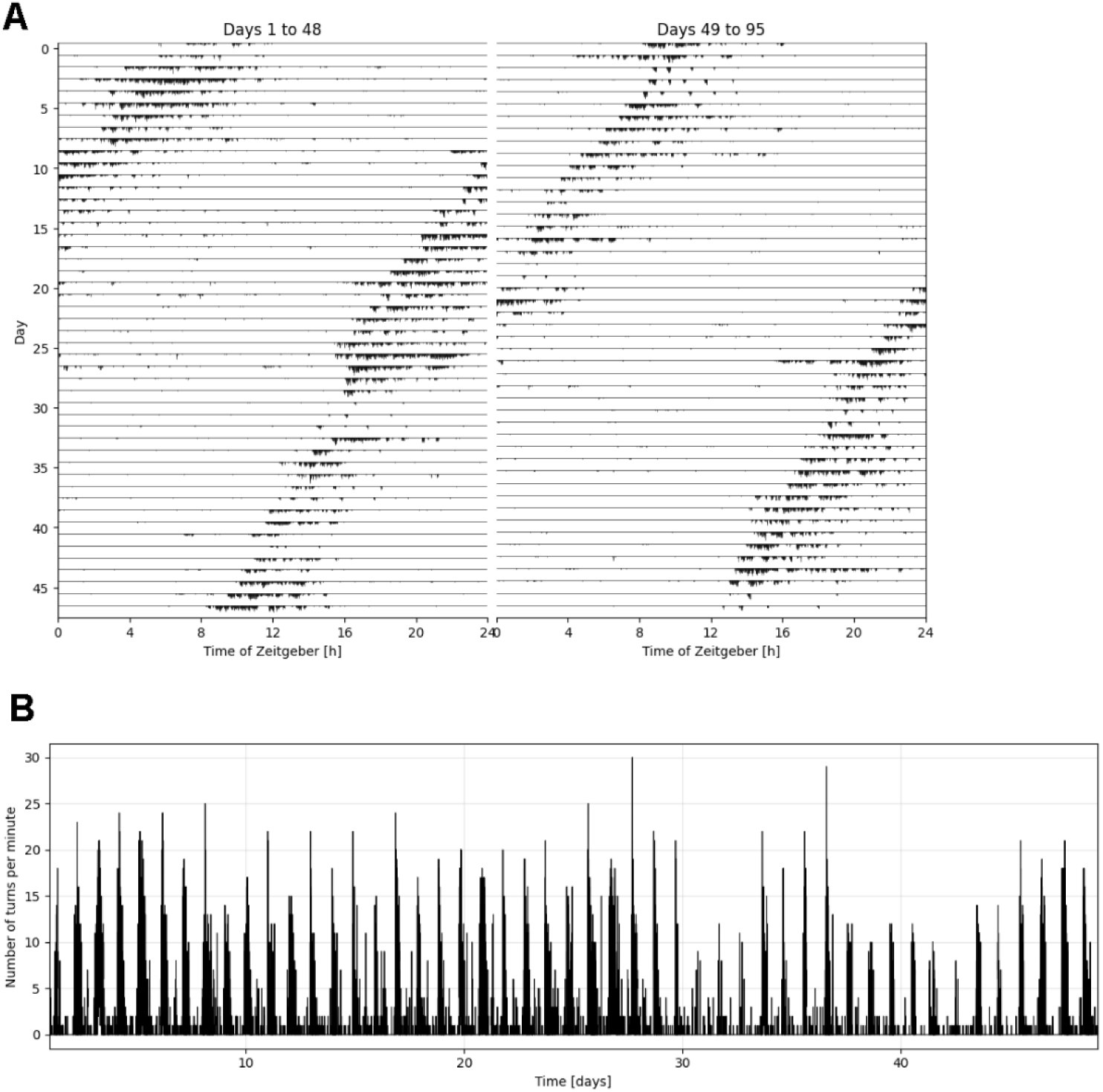
Comparison of continuous activity time series and the corresponding actogram. (A) Actogram showing the activity of a cockroach across the full recording time, split in two panels (days 1-48 and days 49-95). Each row represents one day, with vertical marks denoting activity. (B) Continuous time series of the same activity for the first half of the recording period, corresponding to the left panel in (A).

An alternative way of looking at these behavioral data is as a continuous time series. Fig 1B is a visualization of the data from Fig 1A as such a time series. The height of each vertical bar corresponds to the number of turns per minute. This value will be called activity in the following. Tools from general signal analysis can be used for determining the circadian rhythm, with an excellent review of the available methods given by Refinetti et al. [8]. The resulting frequency plot showing the contribution of a given frequency to the signal is called periodogram. User -friendly software tools are available for setting up both actograms and periodograms (e.g. [9]).

### 1.2 Rhythm analysis

Discrete analyses in the literature focus on determining phase shifts in the circadian rhythm via applied disturbances, such as light pulses at specific Zeitgeber times. These phase shifts can be visualized by kinks in the straight-line connecting activity clusters in the actogram and are indicators for disturbances in the circadian rhythm. Fushing et al. [10] proposed the so-called hierarchical factor algorithm, in which the activity sequence is coded as a digit of base 3. This digit is derived in a hierarchical way. First, recurrence times (times between successive activities) are collected in a histogram, and a threshold probability *α* is defined. Then, a lower bound is defined by the empirical a-quantile *q_α_* and an upper one using the empirical *α*-quantile *q_1-α_*. Recurrence times below *q_α_* are coded with 2, and recurrence times larger than q_1-α_ are coded as 0, with 1 as coding number for the values in between. This procedure can be repeated by omitting the *a-* and *1-a* tails in the histogram, and applying the described coding process to the truncated histogram.

In their analysis, Fushing et al. [11] concentrate on analyzing long rest periods as indicators of the circadian rhythm and its variability. The relation between various signal characteristics and actograms was studied by Sakura et al. [12]. Tackenberg et al. [13] focus on shifts in the circadian rhythm, which may occur after some stimulus. Their main focus is the speed with which the pacemaker resets after a phase-shifting stimulus, as this speed is a fundamental property of circadian clocks. In the paper of Leise et al. [14], both discrete and continuous wavelet analysis are applied to the analysis of activity rhythms. The discrete analysis allows attributing specific frequency bands to the signal, from which the circadian band can be clearly identified. Moreover, it was found that the peak-to-peak phases (variable *DistanceMaxActivity* in the analysis presented below) give the most consistent description of the circadian rhythm. The continuous wavelet analysis, on the other hand, yielded a phase shift between temperature and activity rhythms of hamsters in addition to circadian rhythms.

### 1.3 Current Analysis

All the papers cited above focus on the circadian rhythm and not on ultradian rhythms. As actograms give no direct approach to these rhythms, an approach based on time series will be pursued in this paper with a test series performed with altogether 23 cockroaches kept under constant darkness over more than 40-95 days with the purpose of identifying both circadian and ultradian rhythms.

The data selected represent two groups of cockroaches with different levels of activity. Fig 2 gives typical examples. It can be seen that the maximum number of turns during activity periods stays rather low in the low activity group, whereas more than 5-10 turns per minute were easily reached by the cockroaches in the high activity group. A more quantitative characterization of the two groups will be given in the following.

**Fig 2.**
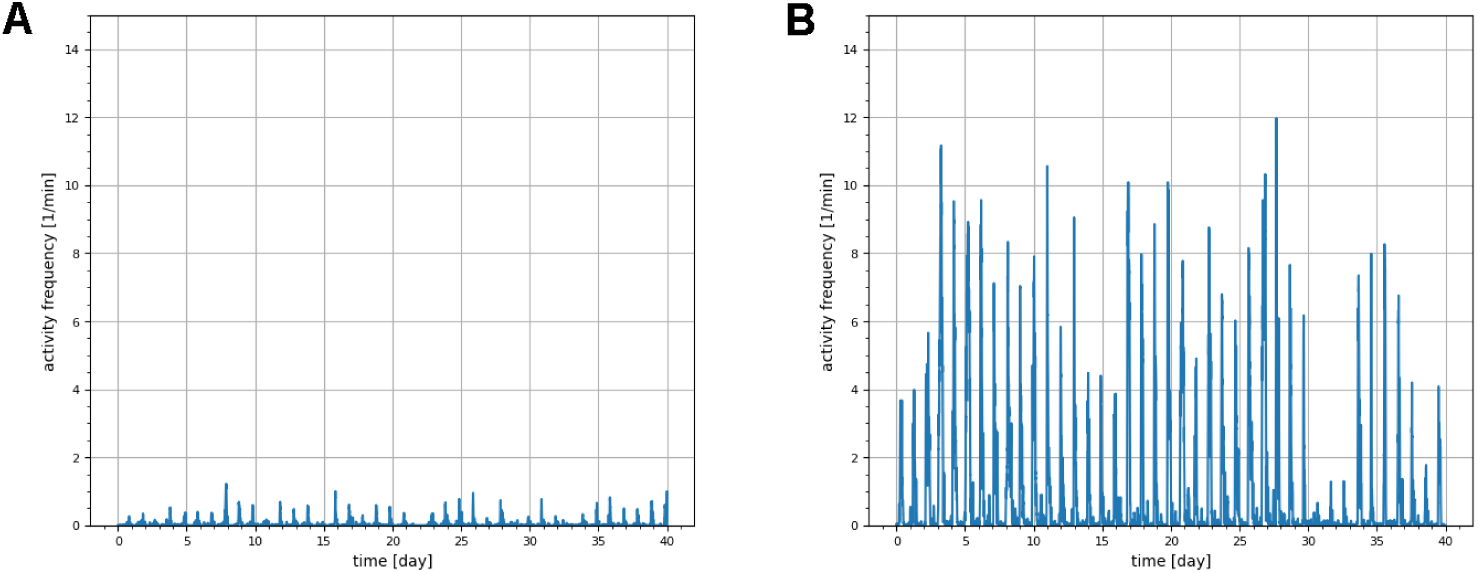
Variability of general level of locomotor activity in the two activity groups. (A) Continuous time series of activity frequency (turns per minute) for a representative animal of the low activity group. It is characterized by consistently low activity with peaks not reaching 2 turns per minute. (B) Continuous time series of activity frequency for a representative animal of the high activity group, showing substantially higher activity, with peaks reaching up to 12 turns per minute.

It is found that there are spontaneous breaks in the activity level, in particular in the high activity group. These breaks can also be found in the low activity group, but are of course less conspicuous in this group. It is highly likely that these breaks are related to experimental conditions (e.g. loss of data in the signal processing unit or obstruction of the wheel), and they have to be considered as outliers in the analysis based on distances between activity impulses.

Unsupervised machine learning [15] will be used to extract temporal patterns from the activity data. This comprises a class of methods that learns patterns, structures, or latent relationships from unlabeled data, meaning no target output variable is provided. The paper is organized as follows: First, metrics of activity will be defined which, in turn, are used to derive a database from the existing activity data. Then the database is split up in two groups of animals, namely a high activity group and a low activity group. Activity rhythms are determined in the following for both activity group. The discussion focusses both on evaluation methods and on features related to ultradian rhythms.

## 2. Metrics of Activity

### 2.1 Defining Time-Distance between Activity

The basic idea of the analysis is given by the fact that the discrete recording of turns per minute can be viewed as being sampled from a continuous process with a sampling rate of 1/60s=0.01667Hz. This has the advantage that established tools from signal analysis can be used for extracting characteristic features from the recordings.

Each turn of the wheel is called activity. The signal noise comes from different sources and is visible at different time scales. The first scale is the individual activity of a cockroach at a given time. The movement may have happened at any instance of the sampling interval of 1 minute, but is recorded at a fixed sampling time. This implies that any recording has an inherent uncertainty of ± 1 minute. Consequently, an elementary activity with one wheel turn has an elementary width of 3 minutes.

The activity is grouped into activity impulses, i.e. groups of consecutive activity. It is assumed that activities impulses can be separated, if they are more than 2 elementary widths apart, leading to an impulse-to- impulse threshold of 9 minutes. A minimal peak height of 0.5 turns is imposed for an activity impulse in order to exclude random disturbances of the wheel turns.

In the preprocessing step, a uniform filter (moving average) was applied to the activity data with width 3 min based on these considerations. A peak analysis was performed using the *find_peaks* method from *scipy. signal* [16] in order to identify activity impulses. This method is solely based on geometrical analysis of the signal curve. The basis is the discrete point cloud obtained by sampling from the filtered signal. Consecutive signal points above a given threshold are merged into one signal section, and the peak is defined as the maximum signal value in that section. Each signal section with a peak has a left root and a right root. Thus, peak analysis converts the signal into activity impulses. Returned variables include the time and the height of the identified peaks *(TimeMaxActivity, MaxActivity),* the left root of the peak *(TimeStartActivity),* and right root of the peak *(TimeEndActivity).* An impulse-to-impulse threshold of 9 minutes was incorporated in the analysis. The area of the activity impulse is estimated by the trapezoid rule and is called *IntensityActivity*.

### 2.2. Determination of activity features

The activity impulses found in the preprocessing step are saved as “activity times” in tuples 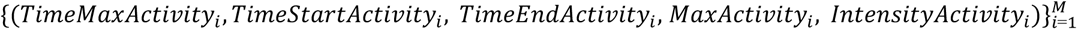, *M* is the total number of identified activity impulses. In this step we calculate features that characterize the activity impulses of the behavior of cockroaches (Fig 3):

**Fig 3.**
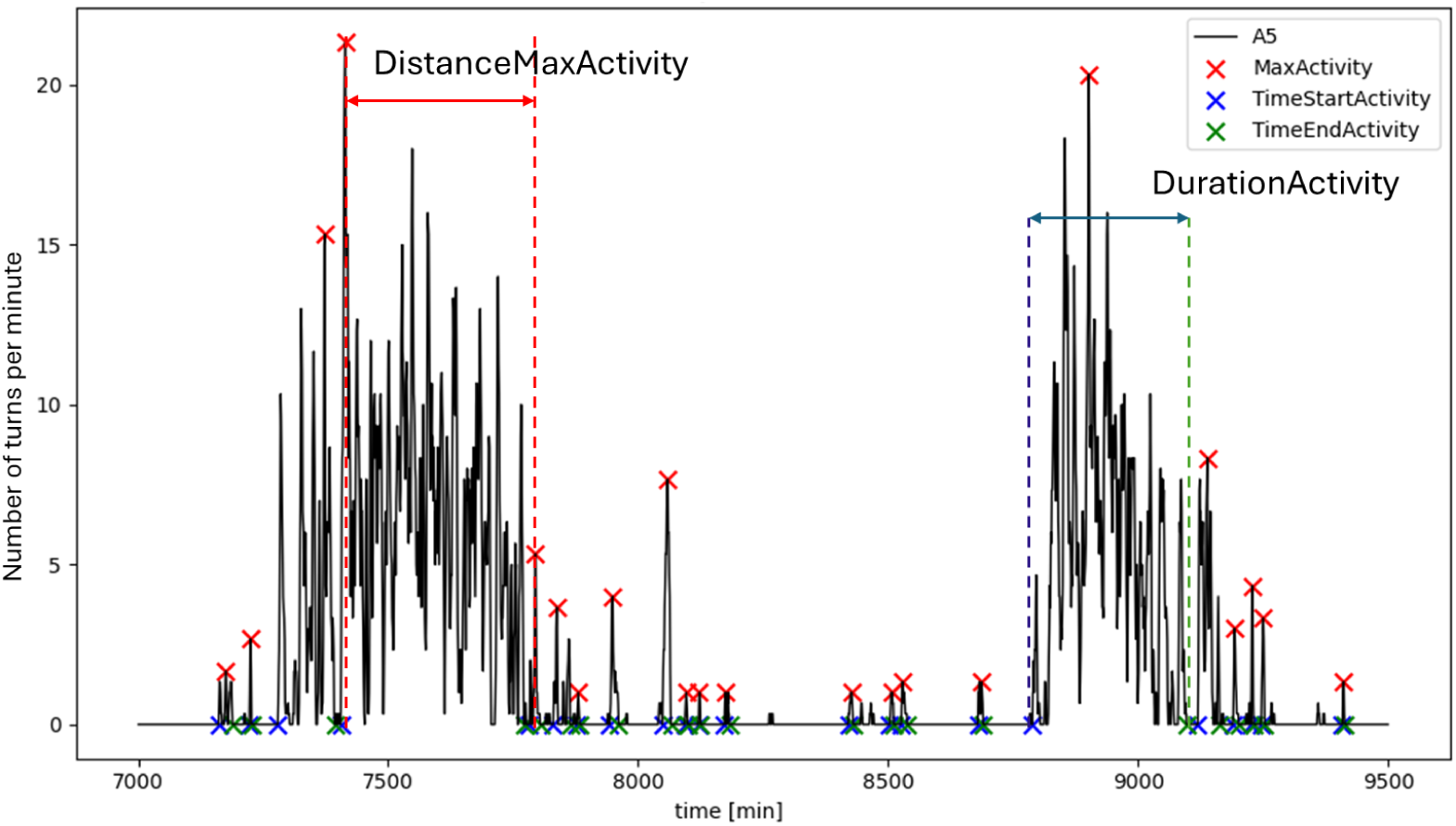
Features for characterizing activity impulses (data from animal A5). Activity impulses were extracted from the filtered activity signal (black line, turns per minute) by peak analysis (*find_peaks, scipy. signal*). Each impulse is characterized by its highest activity, *MaxActivity* (red crosses), occurring at *TimeMaxActivity,* and its onset, *TimeStartActivity* (blue crosses), and offset, *TimeEndActivity* (green crosses). The time between *TimeStartActivity* and *TimeEndActivity* defines the length of an impulse and is denoted by *DurationActivity.* The time between the peaks of two consecutive impulses is denoted by *DistanceMaxActivity*.

We have two variables to describe the **starting** and the **end times of an activity impulse**, *TimeStartActivityi*, *TimeEndActivityi.* With these, we define the **duration of the activity impulse** by:

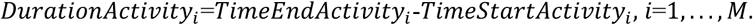

The **duration of the rest period** *i* is calculated as

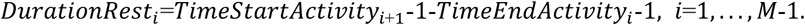

In this setting we can determine the rhythm of cockroach activity in several ways. One way is to calculate the difference of the start times of two subsequent activity impulses:

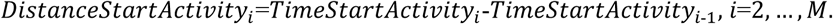

Another possibility is to calculate the time difference of the maxima of two adjacent activity impulses:

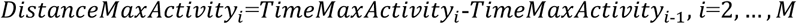

As in the case of activity, we can define a measure for the temporal pattern by the difference of start time of two subsequent rest periods:

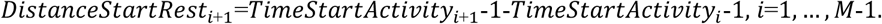

This methodology ensures that each activity period is accurately defined based on the peak analysis facilitating subsequent analyses of activity durations and temporal patterns.

The variables *MaxActivity* and *IntensityActivity* can be used as metrics of activity. The temporal features can be characterized by the distance between the onset of activity impulses *DistanceStartActivity,* the distance between activity impulse maxima *DistanceMaxActivity* and the length of activity periods, *DurationActivity.* Moreover, the variables *DistanceStartRest* and *DurationRest* can be used as metrics of rest.

To analyze the dynamics between active and rest periods in cockroach wheel-turning behavior, we introduce a feature that quantifies the relationship between the duration of an activity impulse (packet) and the subsequent rest period. Given that our algorithm initiates analysis from an activity period, each activity impulse (packet) is inherently followed by a rest period.

We define the **Activity-to-Rest Duration Ratio** for each activity-rest cycle

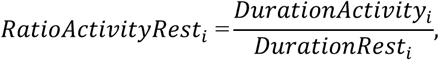

where *DurationResti* is the duration of the rest period immediately following the ith activity impulse for *i* = 1, 2,…, *M-1*.

## 3. Definition of activity groups

### 3.1 Theoretical background

To identify patterns in the cockroach activity data beyond what we can capture with classical rhythm analysis, we employ exploratory, data-driven methods. The goal is to identify groups of individuals that show similar activity patterns. For the following analysis, we focus on the activity metric *DistanceMaxActivity*.

Since the recording durations varied across the animals, the lengths of the vectors *DistanceMaxActivity* differ from each other, and therefore we cannot directly analyze the results obtained from peak analysis. Instead, we transform the *DistanceMaxActivity* values into histograms, which provides a vector of the same length for each cockroach while capturing the distribution of the activity metric [17]. We take an upper bound of 30 hours as larger values of *DistanceMaxActivity* are highly likely to be related to data acquisition problems This interval is divided into several smaller discrete intervals, called bins.

#### 3.1.1 K-Means clustering

For each cockroach we then count how many data points fall into each of those bins. To find an optimal number of bins and, additionally, to find the optimal number of behavioral groups for our cockroaches, we perform a grid search over 3 to 30 bins and 2 to 5 clusters, applying K-Means clustering to each configuration. The clustering quality is then being compared by using two metrics: the Silhouette Score and the Davies-Bouldin Index. This combination is widely used in K-Means clustering validation, as both metrics are highly reliable and they complement each other by evaluating different aspects of cluster quality [18].

The K-Means clustering method is an unsupervised machine learning algorithm [19,20] which groups data points into a predetermined number of clusters, *c,* based on their internal similarity, with the aim of minimizing the distance between each data point and the centroid of its assigned cluster [19]. The algorithm has the aim to minimize the K-Means objective function for cluster centers and memberships [20].

Let *T={t_1_, t_2_,… t_n_}* be a data set in a d-dimensional Euclidian space, *t_t_ e R^d^*, with the distance 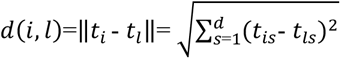, where *t_is_* is the s component of *t_t_* . *A={a1, a_2_,., a_c_*} is the set of *c* cluster centers, and *z_ik_, i=1,*…, *n, k=1, …, c* is a binary variable which indicates if the data point *t_t_* belongs to the *kth* cluster:

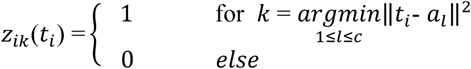

with the centroid *a_k_* following from the following relation:

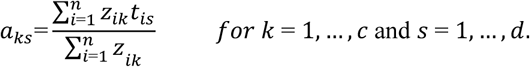

The objective function for cluster centers and memberships

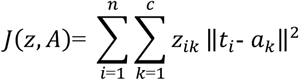

aims at dividing *n* observations into *c* clusters, in which each observation belongs to a cluster *k* with centroid *a_k_*. Minimizing *](z, A)* determines the cluster centroids and the memberships. The minimization is performed using the so-called Expectation-Maximization (EM) algorithm [21].

The quality of the clustering can be measured by the so-called silhouette score. Ideally, a clustering solution should exhibit strong internal cohesion (i.e., similarity within clusters) and clear external separation (i.e., dissimilarity among clusters). The internal cohesion is measured by the intra-cluster distance *δ_k,i_* of point *i* contained in cluster *C_k_,* which is defined as

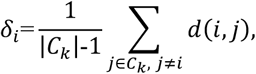

where |*C_k_*| is the number of points in *C_k_* and *d(i,*j)is the Euclidian distance between points *j* and *i* in cluster *C_k_.* For every other cluster *C_t_, I* * *k, 1=1,*.. s, we calculate the average distance of point *i* to all points in this cluster as a measure for separation:

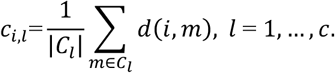

The nearest other cluster is the one with minimum average distance:

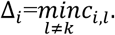

With these quantities, the silhouette value for point *i* can be calculated:

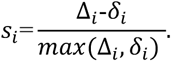

The overall silhouette score *S* then follows from:

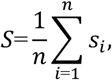

where *n* is the number of data points. The Silhouette score ranges from -1 to 1 with values around 0.5 indicating clear clustering.

The Davies-Bouldin Index quantifies the similarity between clusters by comparing the cluster compactness, which measures the average distance between points in a cluster and their centroid, and the cluster separation, which is the average distance between centroids. Formally, given the same setting as for the Silhouette-Score for a set of *c* clusters with centroids *{a1, a_2_,* …, *a_c_*} and a distance function *d(i, j),* we define for each cluster *C_k_* the cluster compactness by

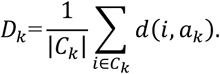

This measures how much the cluster *C_k_* is spread out around its centroid by calculating the average distance of every point *i* e *C_k_* to its centroid *a_k_*. Next, the cluster separation for two clusters *C_k_* and *Cj* is given by the distance of their centroids

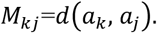

The similarity of two clusters is defined as:

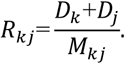

For each cluster, we choose its most similar cluster by

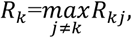

and then we can define the Davies-Bouldin Index as

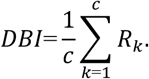

The *DBI* ranges from 0 to infinity, where values near zero indicate well-separated, compact clusters and higher values suggest poor clustering.

Together, the Silhouette-Score and the Davies-Bouldin Index complement each other, as the Silhouette-Score evaluates the cluster quality at the point level, while the Davies-Bouldin Index focuses on the overall cluster compactness and separation.

#### 3.1.2 Self-Organizing Maps

In addition to K-Means, we use Self-Organizing Maps (SOMs) to identify behavioral groups among the cockroaches. Unlike K-Means, SOMs do not assume spherical or well-separated clusters. This makes this method suitable for validating our clustering results and for gaining additional information about the animal population. SOM, founded by Kohonen [22,23], is an unsupervised learning technique used for clustering and to visualize high-dimensional data. This method creates a low-dimensional (typically two-dimensional) grid of neurons, called map, that represents the input data while preserving topological relationships.

As before, we consider a dataset *T={t_1_,* t_2_, .„, t_n_}, where each data point *t_t_ e R^d^* is a d-dimensional feature vector. The SOM consists of *K* neurons, denoted as *K* = {1, . . ., *K},* arranged in a two-dimensional rectangular grid. Each neuron *k e K* is associated with a weight vector *m_k_(j)* e *R^d^* at iteration *j.* At the initial iteration *j* = 0, each weight vector *m_k_(0)* is randomly initialized. During the training phase, SOMs iteratively update these weights by finding the Best Matching Unit 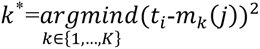, which is the neuron closest to the datapoint *t_i_.* The neurons are updated by:

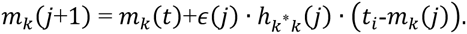

The iteration stops when the difference between the weights becomes small.

### 3.2 Definition of activity groups

#### 3.2.1 Clustering metrics

In Fig 4, we see the Silhouette-Score and Daies-Bouldin Index values of the grid search for our dataset. A number of 3 bins consistently leads to the best results for both metrics. The Silhouette-Score reaches its peak at *c =* 3, while *c =* 2 results in a similarly good score. Meanwhile, the Davies-Bouldin Index reaches its lowest point for *c* = 2 clusters, confirming a number of two clusters to be a good solution. While *c* = 3 also yields a reasonably good Davies-Bouldin Index, it does not reach the same cluster separation as *c* = 2 does. Therefore, we will use *c* = 2 clusters as the optimal solution while *c* = 3 will be analyzed for possible sub-structures within the groups.

**Fig 4.**
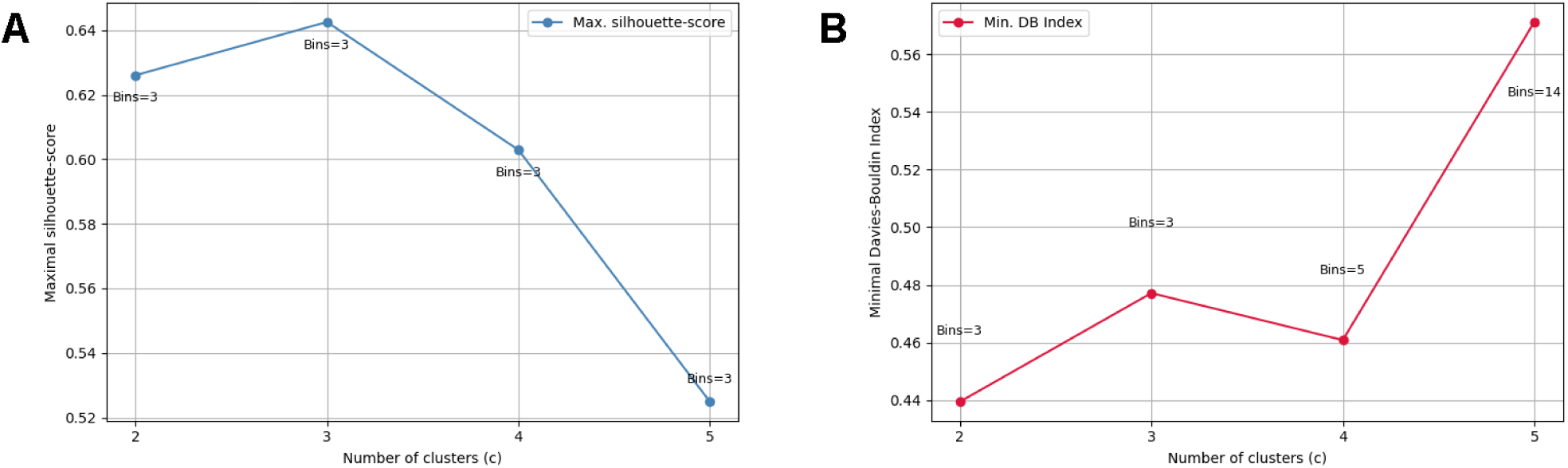
Grid search for optimal number of clusters and bin count. (A) Silhouette score and (B) Davies-Bouldin index for each candidate number of clusters, obtained from a grid search over the number of bins. The corresponding optimal number of bins is annotated at each point. Since a high Silhouette score and a low Davies-Bouldin index indicate good cluster separation, we chose *c* = 2 clusters as the optimal choice. Additionally, *c* = 3 was also examined, as it may provide possible sub-structures.

### 3.3 Results of clustering

As we can see in Fig 5, there are clear differences between the two groups regarding both axes. The high activity group (blue) shows a high normalized density of *DistanceMaxActivity* between 0-10 hours, while having a low density of very long recovery intervals in a range of 20-30 hours. In contrast, the low activity group (orange) shows a noticeably lower proportion of this metric in the 0-10 hours range, accompanied by a higher density of recovery intervals in the 10-20 hours and 20-30 hours bins. This suggests that low activity cockroaches tend to rest longer between two activity impulses than high activity cockroaches do.

**Fig 5.**
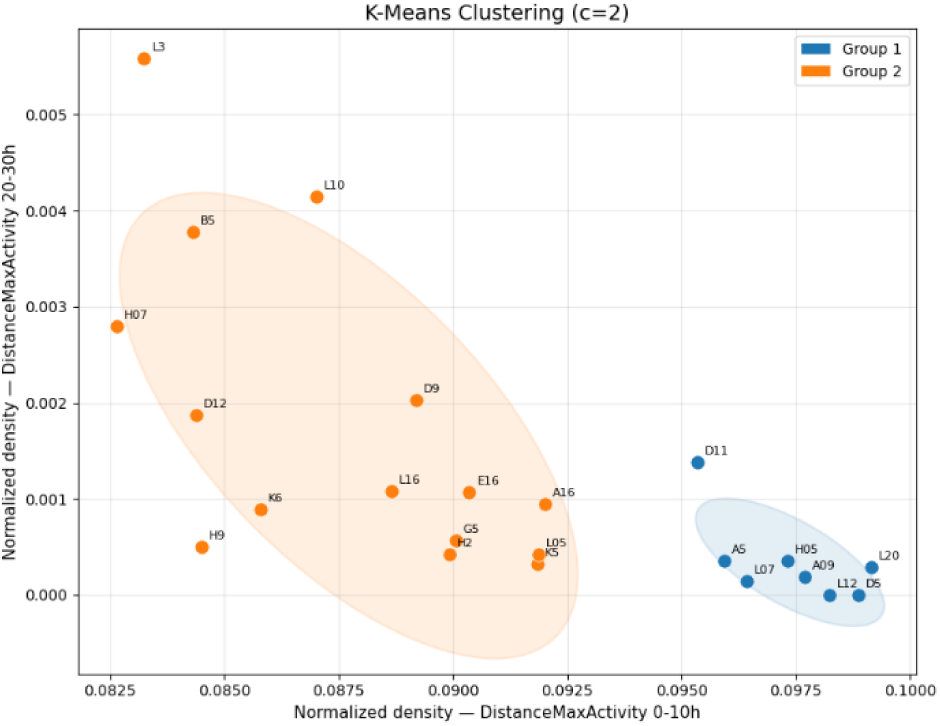
K-Means clustering (*c* = 2) for the 23 cockroaches based on histogram feature vectors of *DistanceMaxActivity* distribution. Each point represents one animal, plotted according to the normalized density of *DistanceMaxActivity* values in the 0-10h bin (x-axis) and the 20-30h bin (y-axis). Two distinct groups are identified by clustering, shown in blue (Group 1) and orange (Group 2).

When slightly varying the parameters, we observed that the animals A16, G5, K5 and L05 are unstable and repeatedly switch their group assignment. They seem to fall between the high activity and the low activity groups, and therefore we will refer to them as border-zone animals. To further investigate whether these animals form a subgroup of their own, we also ran K-Means with 3 bins and *c* = 3 clusters.

Fig 6 confirms that the third cluster (purple) indeed consists of the border zone animals, alongside some other animals from the previous low activity group. This intermediate group shows recovery interval distributions that are neither as short as those of the high activity group nor as long as those of the low activity one. The remaining two clusters correspond to the high activity and low activity groups identified earlier for *c* = 2 clusters. This result for *c* = 3 suggests that the two-cluster solution is already robust, while a third cluster only captures a finer structure within the transition zone between the two main groups rather than introducing a new behavioral type.

**Fig 6.**
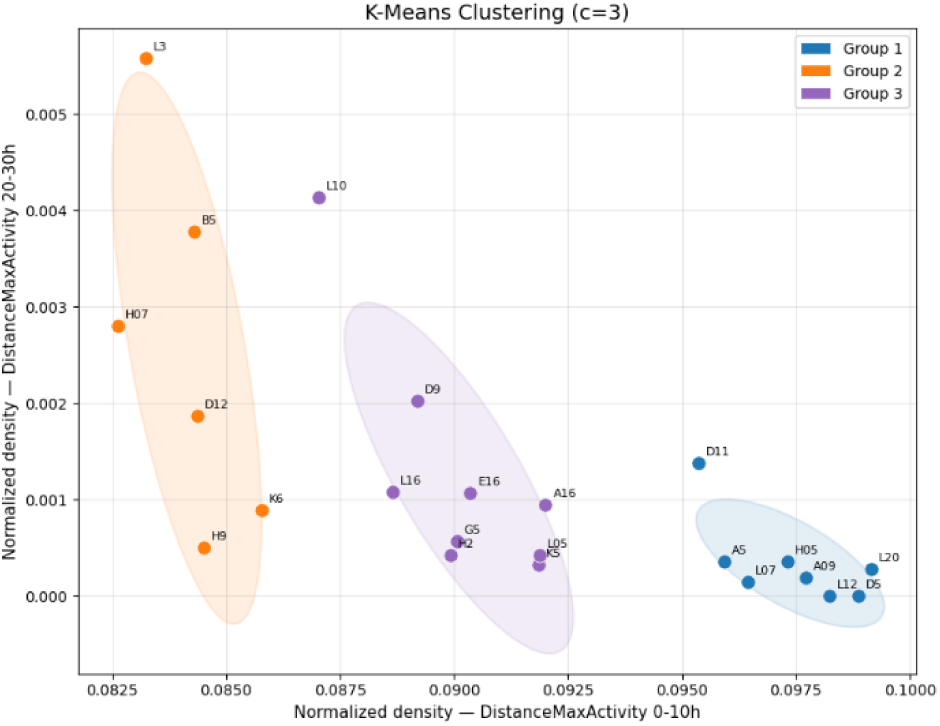
K-Means clustering *(c* = 3) for the 23 cockroaches based on histogram feature vectors of *DistanceMaxActivity* distribution. Each point represents one animal, plotted according to the normalized density of *DistanceMaxActivity* values in the 0-10h bin (x-axis) and the 20-30h bin (y-axis). Three distinct groups are identified by clustering, shown in blue (Group 1), orange (Group 2) and purple (Group 3).

To further validate the clustering results, we applied SOM to this new dataset. Based on the formula *K* ≈ *5√N* introduced by Vesanto and Alhoniemi [24], where *N* denotes the number of data points, the map is determined to have *K* ≈ *5√N* ≈ 24 neurons with *N* = 23 cockroaches, which leads to a 4 x 6 grid. The neighborhood function is adjusted such that approximately half of the map is covered in the initial training phase and the initial learning rate is set to e_0_ = 0.4.

The resulting SOM, depicted in Fig 7, confirms the two-cluster structure with the high activity and low activity animals being clearly separated across the map. The high activity animals are clustered in the upper area of the map, while low activity animals are positioned at the bottom. The groups are separated by a dark region that reflects high distances between the neuron weights and marks a clear boundary between the two clusters. Within this zone lie the previously mentioned border zone animals A16, K5 and L05 alongside the previously stable high activity animals A5 and L07 as well as the low activity animals D9 and E16.

**Fig 7.**
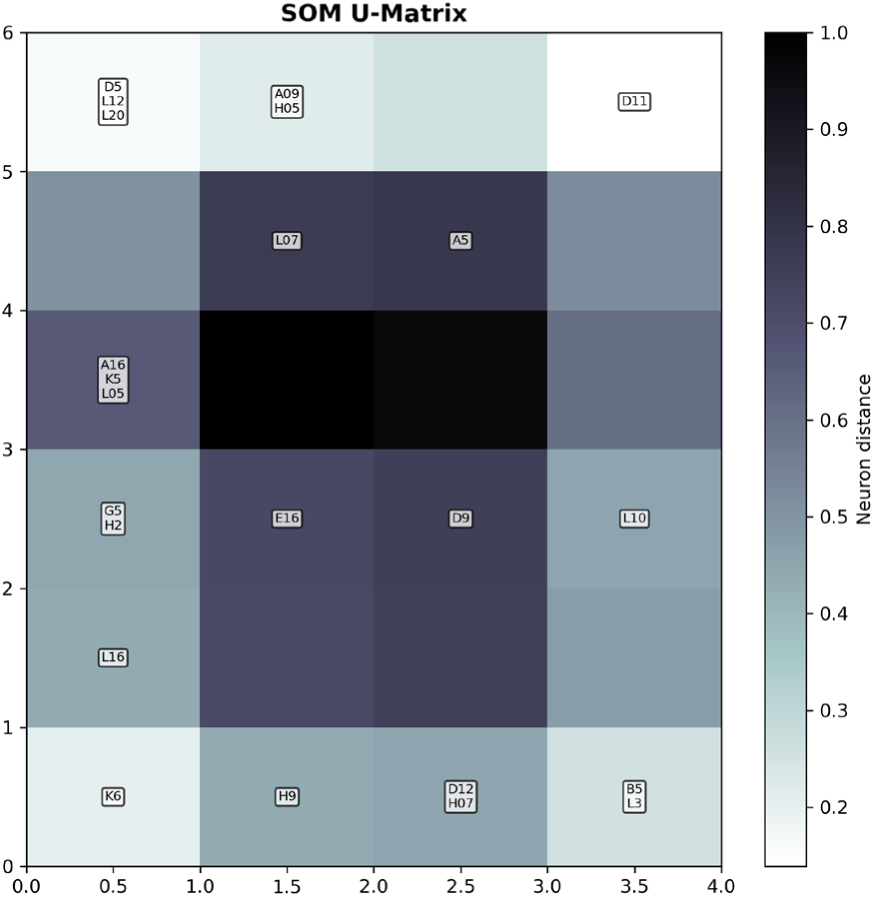
U-matrix representation of the SOM showing activity-based groups of cockroaches. Dark regions indicate large neuron distances and mark boundaries between groups. Labels show the animals mapped to each neuron. The light area at the top corresponds to high activity animals, while the bottom area shows low activity ones.

Comparing the animals placed in the darker region to those identified as border zone animals through K-Means clustering shows that the two groups are not identical. In contrast to the *c* = 3 clustering, where the intermediate group consisted only of low activity cockroaches, the SOM places two high activity ones alongside the border zone animals. This suggests that the dark region primarily reflects proximity to the cluster boundary rather than a third subgroup and that the SOM overall supports the two-group structure identified through K-Means clustering.

## 4. Analysis of Rhythms

### 4.1 Temporal patterns

A straightforward visualization of temporal patterns can be done by plotting the activity of the animals on a scale of several days. Thus, the circadian rhythm is clearly visible (see Fig 2). Ultradian rhythms, however, are more difficult to visualize as they vary on a daily level. Therefore, some kind of scaling is needed which cuts the signal train in slices corresponding to one circadian period. Averaging the activity in these slices, in turn, will give a database for visualizing and understanding the activity patterns within one circadian period. This implies that we first have to extract circadian rhythms, before looking for temporal patterns on a smaller time scale. In the analysis presented here, the circadian patterns are extracted using /2 periodograms.

#### 4.1.1 χ^2^ periodogram

This method is particularly suitable for data which are not evenly spaced as it is the case for the activity data of cockroaches. The basic idea is to select a given period *P* and determine tuples 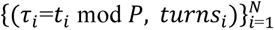. The interval [0, *P]* is then divided into *K* bins and the number *Oj* of counts in bin *bj* is determined from:

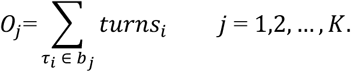

If the period is not relevant for the time series under consideration (null hypothesis), then the number of counts per bin should be uniformly distributed with an expected number *Ej =K/N* of counts per bin. The quantity

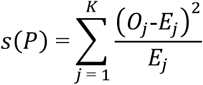

is approximately χ^2^-distributed with *K-1* degrees of freedom. Values *s(P)* much larger than the expected value *K-1* indicate that the null hypothesis has to be rejected. In other words, *P* is contained in the frequency spectrum of the data.

Fig 8 shows the typical periodograms obtained for the high activity and the low activity groups. Periods in the range between 1 hour and 25 hours are taken into consideration, and the number of bins was chosen to be *M=50.* Different values of *M* did not have a strong impact on the overall character of the periodograms.

**Fig 8.**
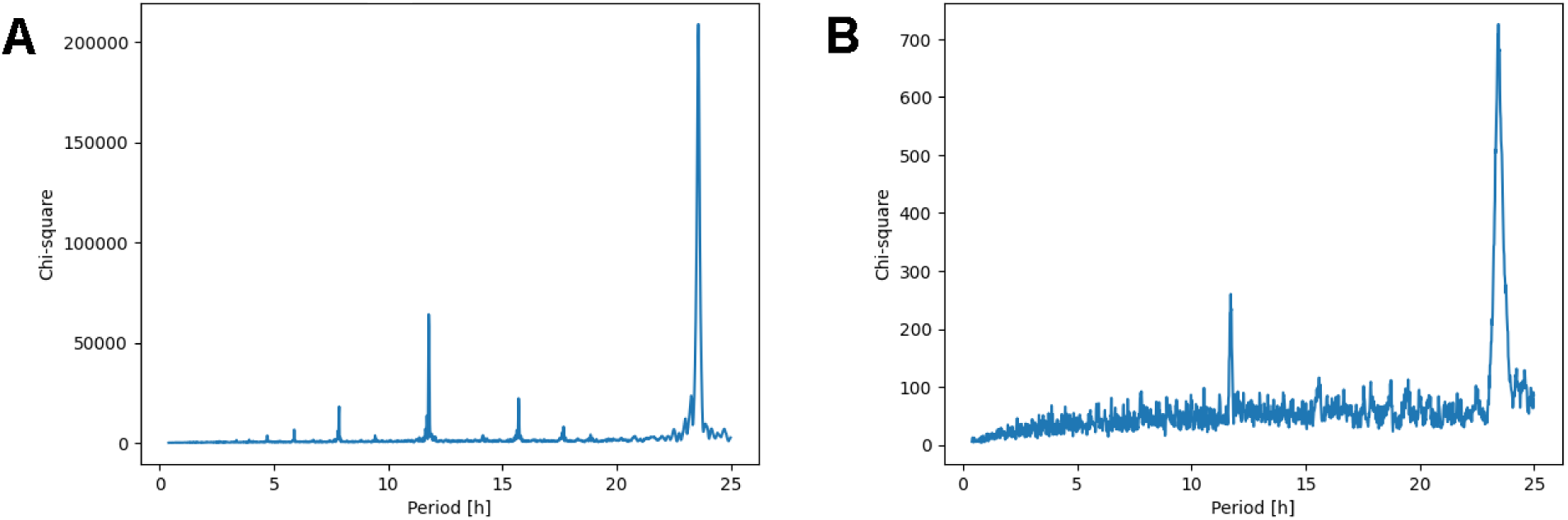
Chi-square periodogram analysis for periods between 1 hour and 25 hours. (A) Representative animal from the high activity group, showing a dominant peak near 24h, corresponding to its circadian period. (B) Representative animal from the low activity group, showing a comparatively smaller peak near 24h. Note that the y-axis scales differ substantially between the panels (0-200.000 in A vs. 0-700 in B), reflecting the overall lower number of turns in the low activity group.

Fig 8 shows that the circadian rhythm has some variability, but is clearly found in both groups at about 23.5 hours. Moreover, the second harmonic is clearly visible with a period of half the circadian, even though the signal is less clear in the low activity group. The higher amount of noise in the periodogram of the low activity group is related to the reduced total number of turns in the low activity group which amounts to about one tenth of that of the high activity group. The analysis of the high activity group in Fig. 8A suggests that there are other characteristic periods. However, these correspond to higher harmonics and the signal asymmetry.

#### 4.1.2 Scaling

Once the circadian period *T_c_* is known for each animal, scaling can be implemented. A starting point is defined by searching for the highest value of the *MaxActivity* value within the first few hours of the recording. It was found that a starting point defined by searching over the first 10 hours gave a fairly robust starting point for almost all animals. Then the signal train is divided into slices of length *T_c_* beginning at the starting point, and the corresponding characteristic times are normalized by calculating *T_i_=TimeMaxActivity_i_ /T_c_*. Fig 9 illustrates the slicing procedure, where the blue crosses indicate the slicing intervals. It can be concluded that the nightly activity bouts are always at the boundary of the slicing interval, and that activities i and *j* with the same value of the normalized time *t_i_= t_j_* are very likely to have occurred at the same phase shift with respect to the circadian period.

**Fig 9.**
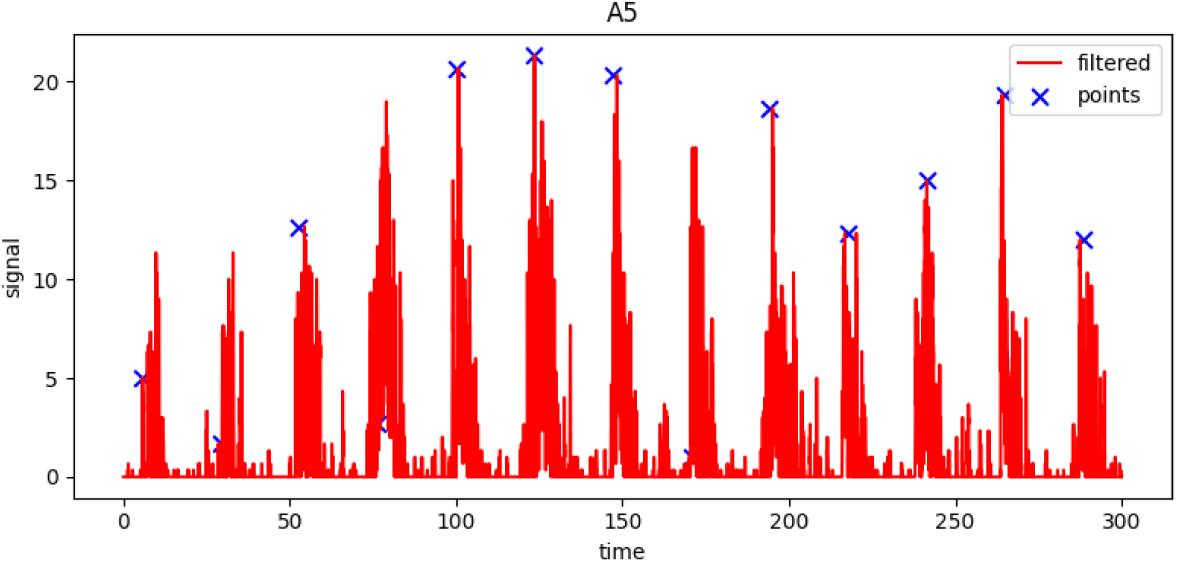
Example of the slicing procedure used for scaling activity data. The filtered activity data (red line) of the representative animal A5 is divided into slices of length *T_c_* (the animal’s circadian period). Blue crosses mark the starting point of each slice, with the first one marking the highest *MaxActivity* value within the first 10 hours of recording, and each following slice begins exactly *T_c_* after the previous one.

Then a regular grid is defined with *t_k_, k =* 1,., *K i_k_e* [0, 1], and *d_t_=DistanceMaxActivityiIT_C_, I* =1, …,*L, d_t_* e [0, *d_max_].* The cutoff value *d_max_* for the scaled activity metric *d_t_* was chosen to be one, as larger time intervals with no activity are very likely caused by data loss. We can define the number of hits *n_lk_* at *t_k_, k=1, …, K* and *DistanceMaxActivityi, 1=1,*…, *L:*

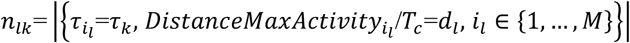

and plot the tuples *(r_k_, d_t_,n_lk_)* in a heatmap in order to visualize the distribution of the activity metric within the circadian cycle.

#### 4.1.3 Logarithmic heatmaps

Heatmaps proved to be rather non-informative, as most of the values of the normalized activity metric *d_t_* were rather small and there was no visible spread of the data in the parameters space. Taking *d_t_* = *log_e_ d_t_* improved the spread and hence the information content of the heatmaps. The results are shown in Fig 10 and Fig 11.

**Fig 10.**
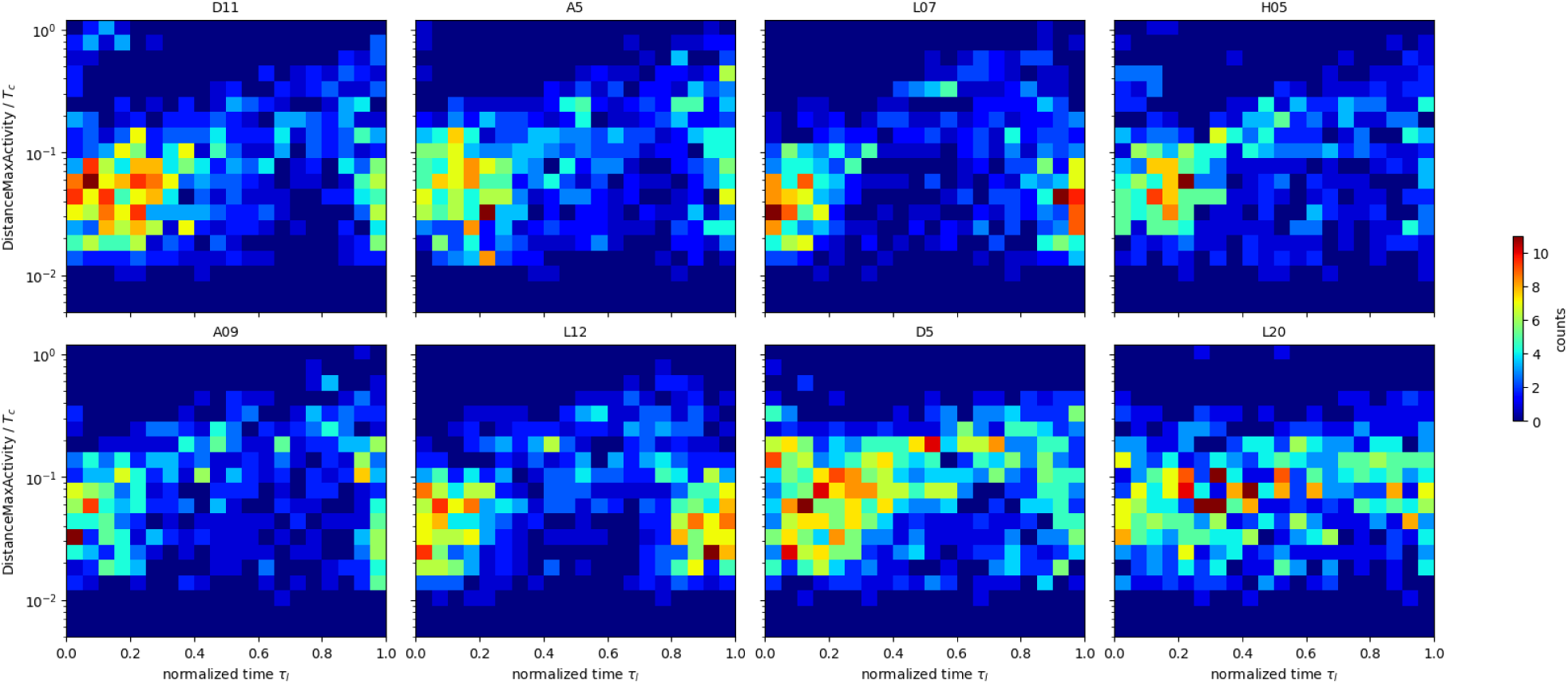
Logarithmic heatmaps for each animal in the high activity group. These heatmaps visualize the distribution of activity within the circadian cycle for each cockroach of the high activity group. The normalized time *τ_i_* (x-axis) is plotted against the log-transformed activity metric *d_l_* (y-axis) and the colors indicate the number of activity impulses.

**Fig 11.**
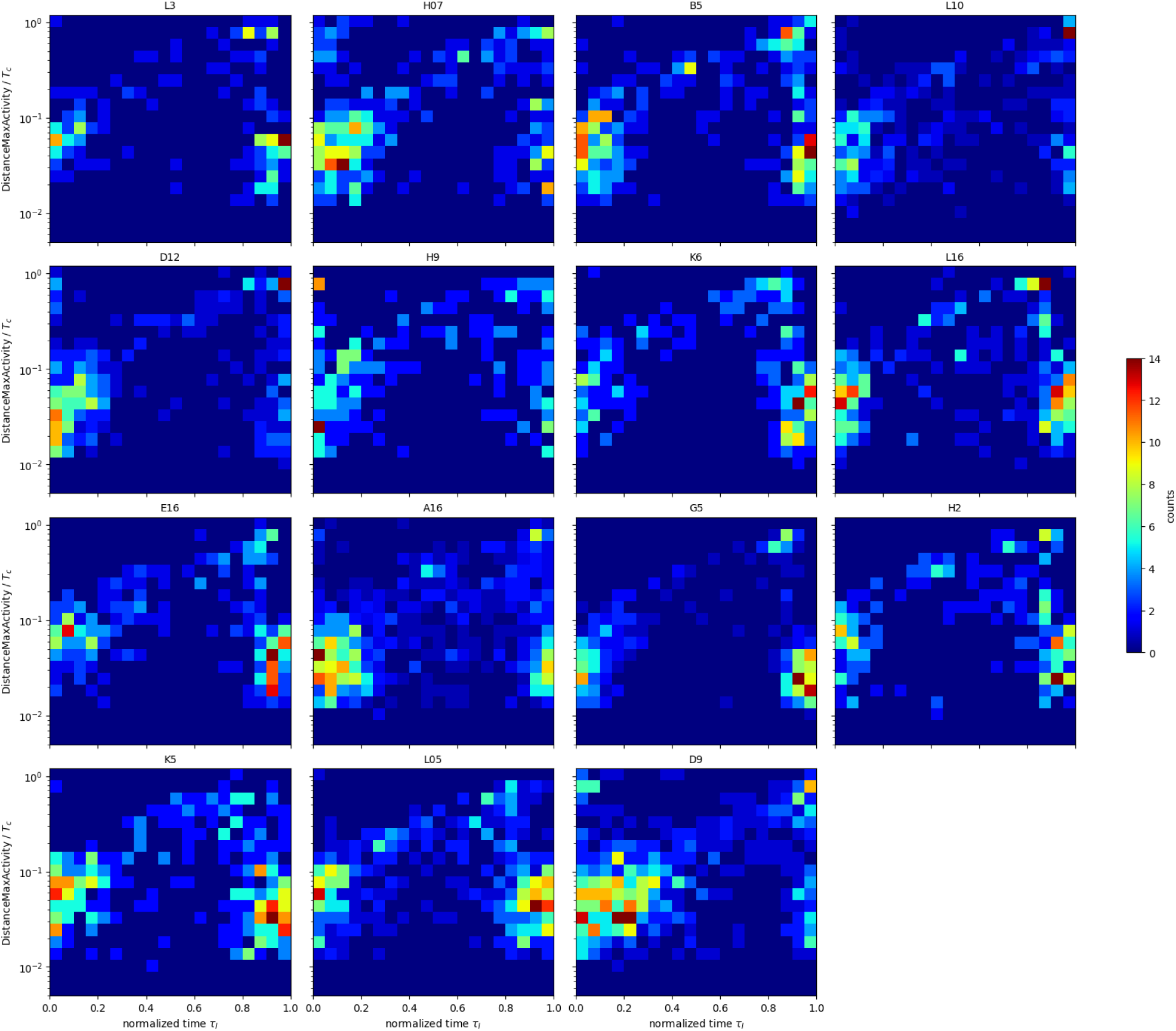
Logarithmic heatmaps for each animal in the low activity group. These heatmaps visualize the distribution of activity within the circadian cycle for each cockroach of the low activity group. The normalized time *τ_i_* (x-axis) is plotted against the log-transformed activity metric *d_l_* (y-axis) and the colors indicate the number of activity impulses.

In both groups, the periodic activity bouts are clearly visible as a cluster located near the lower left or lower right boundary of the grid. In the areas with *r_k_* ≈ 0.5, few activity peaks have been found and the relative activity metric *d_t_* attains higher values.

The differences between the high activity and low activity groups are hard to judge, as the differences between the individual animals are quite significant. However, the picture becomes much clearer, if we sum over all animals:

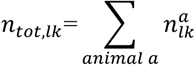

and plot *(r_k_, Hi, n_totAk_)* in a heatmap for each activity group. Fig 12 shows the results. When comparing these diagrams, it has to be kept in mind that the number of animals in the low activity group is higher, and that hence the numbers *n_tot,lk_* cannot be compared in their absolute values.

**Fig 12.**
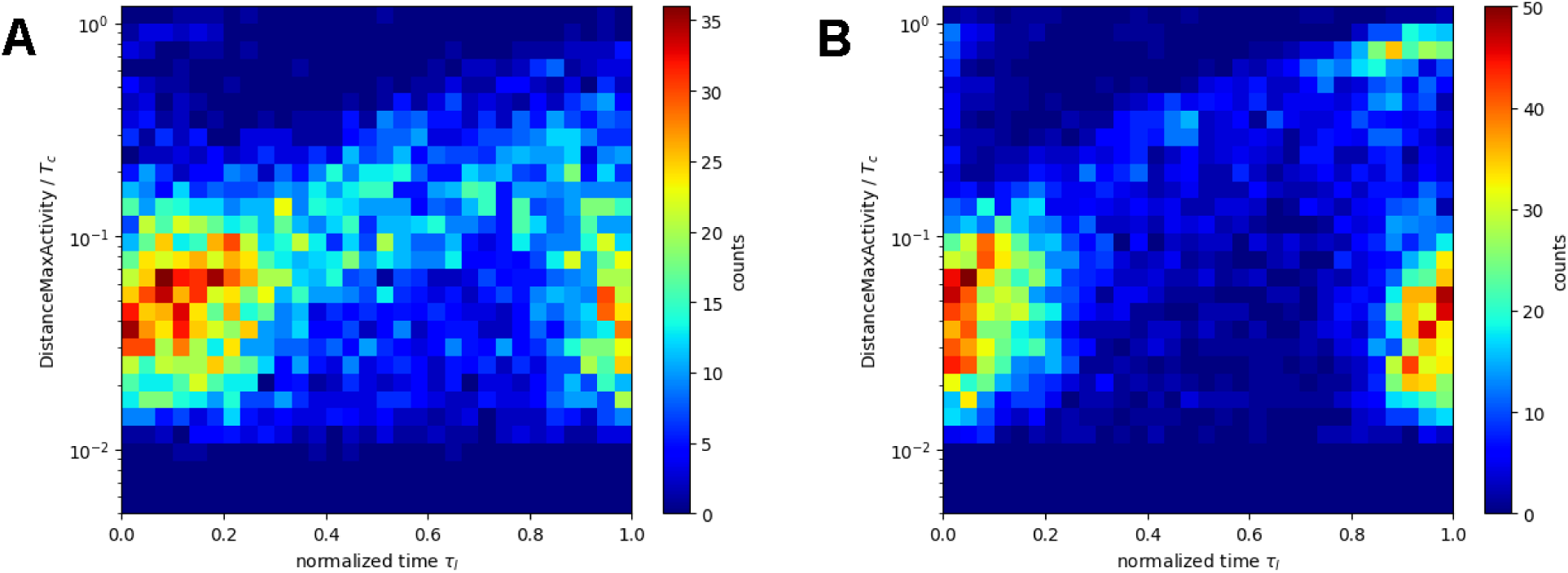
Average logarithmic heatmaps for both activity groups. Heatmaps showing the distribution of the activity metric within the circadian cycle, averaged across all animals within each group. (A) High activity group. (B) Low activity group.

Both the high activity group and the low activity group have a pronounced cluster of values with *d_t_* ≈ 0.05, *t_k_* >0.9 or T_fc_<0.2. These clusters correspond to the circadian activity bursts., We obtain *DistanceMaxActivityl* ≈ 1 *hour* when going back to the original value of *DistanceMaxActivityl=T_C_* • *d_t_* with an average value of *T_c_.* This means that activity peaks occur about every hour during periods of increased activity. The high activity group shows much more activity in the intermediate range between the circadian bursts *0.3<r_k_*<0.8. The heatmaps indicate the typical value of *d_t_* lies around 0.2 for the high activity group, i.e. *DistanceMaxActivityl ∼* 5 *hours* indicating that rest times between bursts of activity are much longer in this period. The probability of having an activity burst is reduced for the low activity group compared to the high activity group, but those animals having bursts of activity during this period do so with similar characteristics. The most striking difference between the low activity group and the high activity group is that there is a clear cluster for the low activity group at *d_t_* ≈ 0, i.e. *DistanceMaxActivityl ∼ Tc-*This implies that there are days with vanishing activity between the circadian bursts for the low activity group.

### 4.2 Gaussian Mixture Model

The Gaussian mixture model can be applied to the data, once a peak analysis has been performed beforehand, and the metrics described in Section 2 have been determined. As the basis hypothesis of the analysis is that the signal is quasi-continuous, because the cockroach could turn the wheel at any instant, and is discretized by the sampling rate and the experimental procedure (only full turns are counted), continuous probability distributions can be employed for describing the variability of the signal. In a first step, experimental data for *DistanceMaxActivity,* were collected in histograms and compared with each other.

Fig 13 shows the result obtained with data containing the low activity and the high activity groups. It has to be kept in mind that the appearance of a histogram depends strongly on the bin size, and the standard bin size chosen in Fig 13 does not agree with the optimal bin size proposed in [25]. However, no quantitative evaluations will be made on the basis of these histograms, and a fairly coarse binning was chosen for the ease of visualization. It can be seen the activity metric has a heavily skewed density functions with rather long tails in both cases. Some difference in in the shape of the density seem to exist, as there is presumably a longer tail for the low activity group.

**Fig 13.**
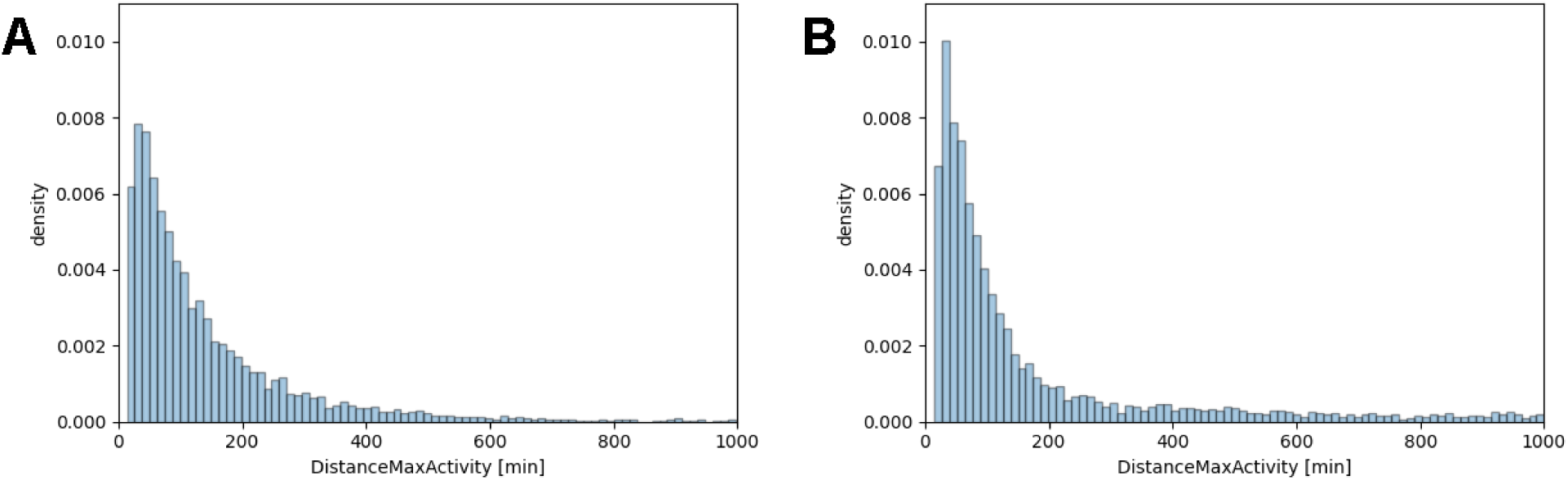
Histograms of the activity metric *DistanceMaxActivity*. (A) High activity group. (B) Low activity group.

A skew-symmetric distribution family with a long tail is the lognormal distribution. However, a preliminary analysis showed that the decrease of the lognormal probability density for values larger than the mode is too fast and does not represent the probability mass contained in the sample interval around one to two hours. Therefore, a multi-modal statistical model was used based on a Gaussian mixture [26] of the logarithmic random variables with the following probability density:

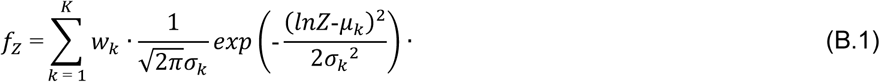

where *Z* denotes the random variable, and the weighting factors are related by w1+… *+w_K_ =* 1.

Estimates of the parameters of the corresponding cumulative distribution function were obtained using *GaussianMixture* from *sklearn.mixture* which uses an EM [21] (expectation maximization) algorithm as mentioned above.

The number *K* of components in the Gaussian mixture model has to be determined separately. For this purpose, some criterion penalizing a very high number of parameters is needed, as increasing the number of components increases the number of fitting parameters which will always improve the match between the data and the estimated distribution.

AIC (Akaike Information Criterion, [27]) and BIC (Bayesian Information Criterion [28]) are popular penalizing criteria in this context. AIC is derived as an estimator of the expected Kullback-Leibler divergence between the unknown data generating density *g(Z)* and the fitted model *f_z_(Z; θ)* where *θ* denotes the corresponding parameter vector. We have

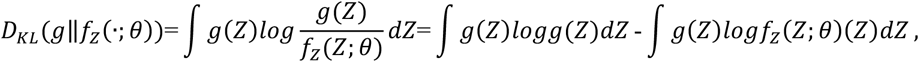

where *D_KL_ (g‖f_z_*) is the standard notation for the Kullback-Leibler divergence. As the first term on the right- hand side does not depend on the parameters *0,* minimizing *D_KL_(g‖f_z_(•;*θ)) is equivalent to maximizing the second term which corresponds to the expectation value of the log-likelihood function *l(θ).* The maximized log-likelihood function with parameters θ̂ is optimistic in the sense that it gives large values for a higher number of parameters. Therefore, we have to penalize by the number *p* of parameters. The final relation for AIC is given by:

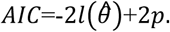

This implies that models with a lower value of AIC are better.

The BIC value, on the other hand, starts from the marginal likelihood, where *n(0)* is the prior distribution of the parameters.

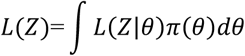

Using log-likelihood instead of likelihood leads to

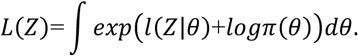

Expanding the exponent in the integrand around its maximum θ̂ up to second order and taking the logarithm on both sides leads to:

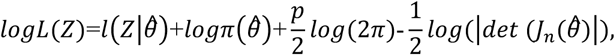

where the last two terms are related to the Lagrange remainder in the Taylor expansion, *p* is the dimension of the 0-space as defined above and *n* is the number of data points (i.e. the dimension of the Z-space). The term *detlJ_n_(6)l* can be shown to be of the order *n^p^.* We get from this:

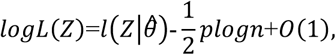

and, finally:

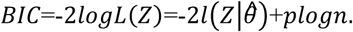

Comparison of AIC and BIC shows that BIC penalizes stronger for large datasets and therefore tends to support smaller models.

Fig 14A and Fig 14B show the AIC and BIC-values found by fitting a Gaussian mixture model to the logarithmic metric *logDistanceMaxActivity.* It can be seen that the BIC criterion favors a three-dimensional model for the high activity group, whereas a five-dimensional model should be employed in the low activity case. However, four components give good values for AIC and BIC in both cases. Now, selecting mixture models with the same dimension facilitates comparison significantly, and it was therefore decided to rely on a four-component Gaussian mixture model for the final evaluation. With that, the data can be fitted very well as shown in Fig 14C and Fig 14D. The parameters obtained are summarized in Table 1. It can be seen that the parameters and the weights of the first two components in the Gaussian mixture model are very similar in the two activity groups, whereas components #3 and #4 seem to be different.

**Fig 14.**
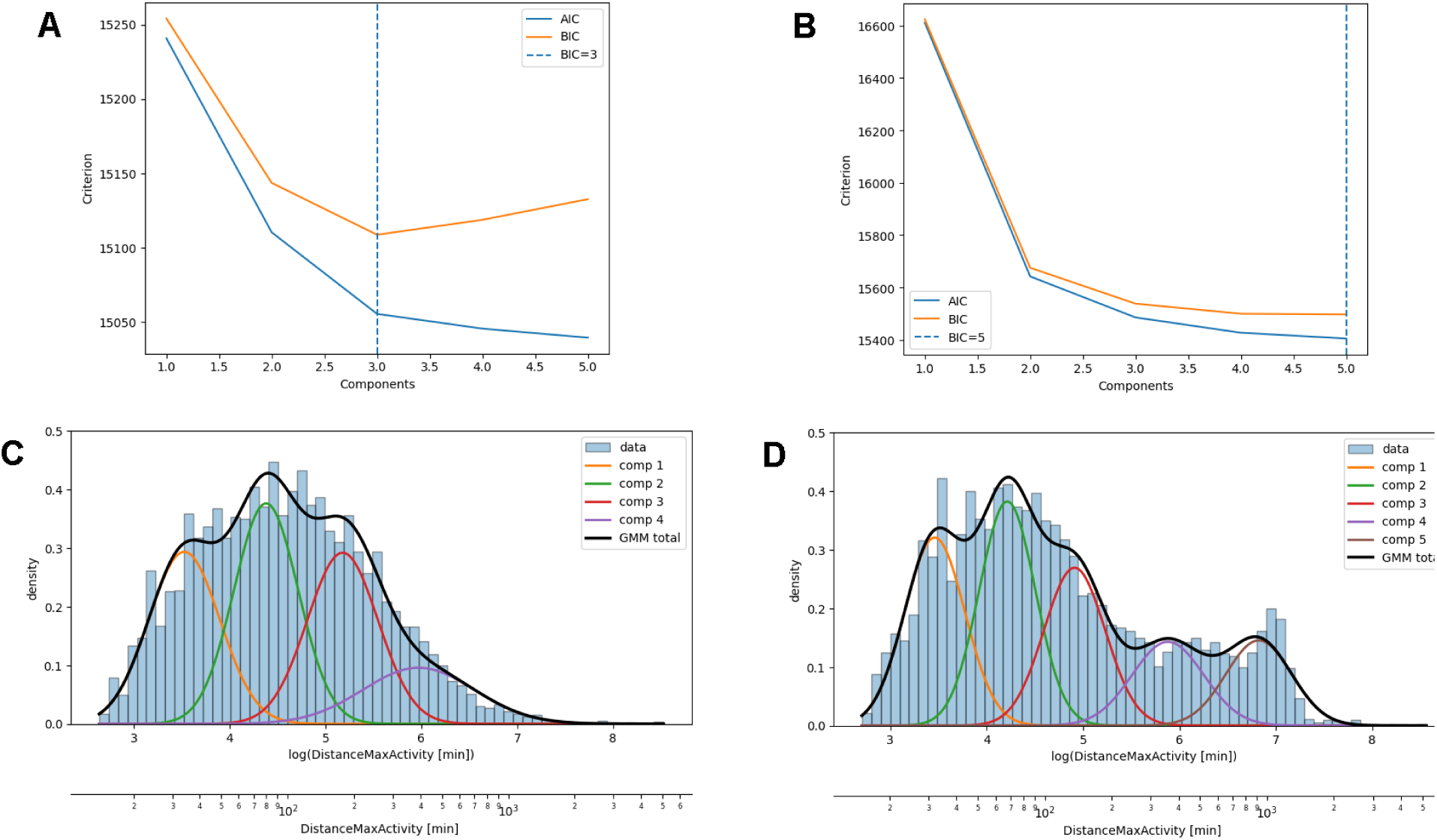
Gaussian mixture model for the metric *DistanceMaxActivity*. (A,B) Penalizing metrics (AIC, BIC) for the Gaussian mixture model fitted to log-transformed *DistanceMaxActivity* as a function of the number of components, for high activity group (A) and the low activity group (B). The dashed lines mark the number of components favored by the BIC. (C,D) Histograms of log-transformed *DistanceMaxActivity* with the fitted Gaussian mixture model for (C) the high activity group with 4 components and (D) the low activity group with 5 components. The black curves show the sum of the individual components in each case.

**Table 1.** Parameters of the Gaussian mixture model.

|  | High activity group |  |  | Low activity group |  |  |
| --- | --- | --- | --- | --- | --- | --- |
| compo-<br>nent | weight | $E[X], [h]$ | $\sqrt{Var[X]}, [h]$ | weight | $E[X], [h]$ | $\sqrt{Var[X]}, [h]$ |
| 1 | 0.274 | 0.607 | 0.234 | 0.307 | 0.624 | 0.232 |
| 2 | 0.317 | 1.41 | 0.487 | 0.347 | 1.52 | 0.565 |
| 3 | 0.270 | 3.17 | 1.210 | 0.177 | 4.12 | 1.85 |
| 4 | 0.140 | 7.74 | 4.910 | 0.169 | 14.2 | 6.36 |

## 5. Discussion

### 5.1. Computational Complexity

Uniform filtering has complexity *O(N),* whereas the complexity of the *find_peaks* is somewhat higher with *O(NlogN)* as a certain amount of searching is necessary to determine local maxima. Hence, a complexity of *O(NlogN)* is attained in summary in the preprocessing step. Straightforward computation of periodograms has complexity *O(p* • *N),* where *p* is the number of periods taken into consideration. The complexity of clustering using K-Means is *O(k* • *N)* per iteration. Gaussian mixture using EM iteration has complexity *O(m* • *N)* where *m* is the number of components with an additional *O(N)* related to taking logarithms.

### 5.2 Algorithms

The machine learning algorithms employed are k-means clustering and SOM for dividing the cockroach population into groups of different activity level, and a Gaussian Mixture Model for finding temporal patterns in the activity metrics.

#### 5.2.1 Clustering

At first glance, classical statistics might seem more suitable as unsupervised learning methods such as SOM and K-Means, as results obtained with statistics are normally intuitively more appealing. However, it has to be kept in mind that most statistical tests are based on some fundamental assumptions on the underlying distributions such as homogeneity or near normality, which are clearly violated in the current database.

An example of the performance of a statistics-based analysis is given in the following. High activity animals should have larger values of the metric *DurationActivity* and smaller values of the metric *DurationRest.* Consequently, the mean value of the metric *RatioActivityRest* seems to be suitable for characterizing the activity level. Comparing the results obtained for the previously identified high activity and low activity groups shows that some animals with rather large values of the *RatioActivityRest* variable have been placed in the low activity group and vice versa (see Fig 15). However, clustering based on a threshold of the mean value of *RatioActivityRest* equal to one lead to less clear effects in the rhythm analysis. The tail of the histogram used for the Gaussian Mixture Model was less prominent for the low activity group, and the cluster at *DistanceMaxActivity* ≈ *T_c_* almost disappeared. This indicates that defining low or high activity on a single statistical moment is less reliable than the methods taken from machine learning. A second trial was made by comparing distributions using Mann-Whitney test. No meaningful clustering could be obtained as translational invariance of the test could not be achieved, i.e. if animal a1 is compatible with animal a2, and a2 with a3, it is not guaranteed that a1 is compatible with a3.

**Fig 15.**
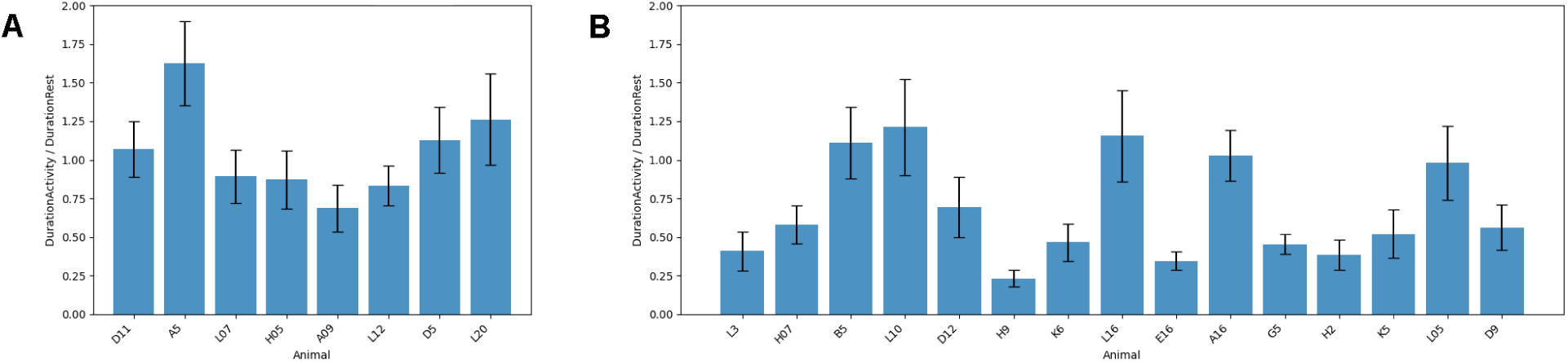
Mean values of *RatioActivityRest* including 95% confidence intervals. (A) High activity group. (B) Low activity group. Each bar represents one animal.

#### 5.2.2 Activity metrics

The analysis presented in the paper is mainly based on the metric *DistanceMaxActivity.* Alternative metrics such as *DistanceStartActivity* or any other of the metrices proposed in Section 2 could have been used instead, and it is not self-evident that *DistanceMaxActivity* is more suitable than the others, and that the findings presented here are in a sense invariant of the metric. Therefore, the evaluations presented in Section 4 were repeated using the metrics *DistanceStartActivity* and *DistanceStartRest.* It was found that there was virtually no change in the results if *DistanceStartActivity* is used, whereas *DistanceStartRest* showed a larger scatter and less structure in the histogram, see Fig 16. However, the clusters in the heatmap and the components of the Gaussian mixture model are quite similar. Hence, the implications of the rhythm analysis do not depend on the choice of the activity metric.

**Fig 16.**
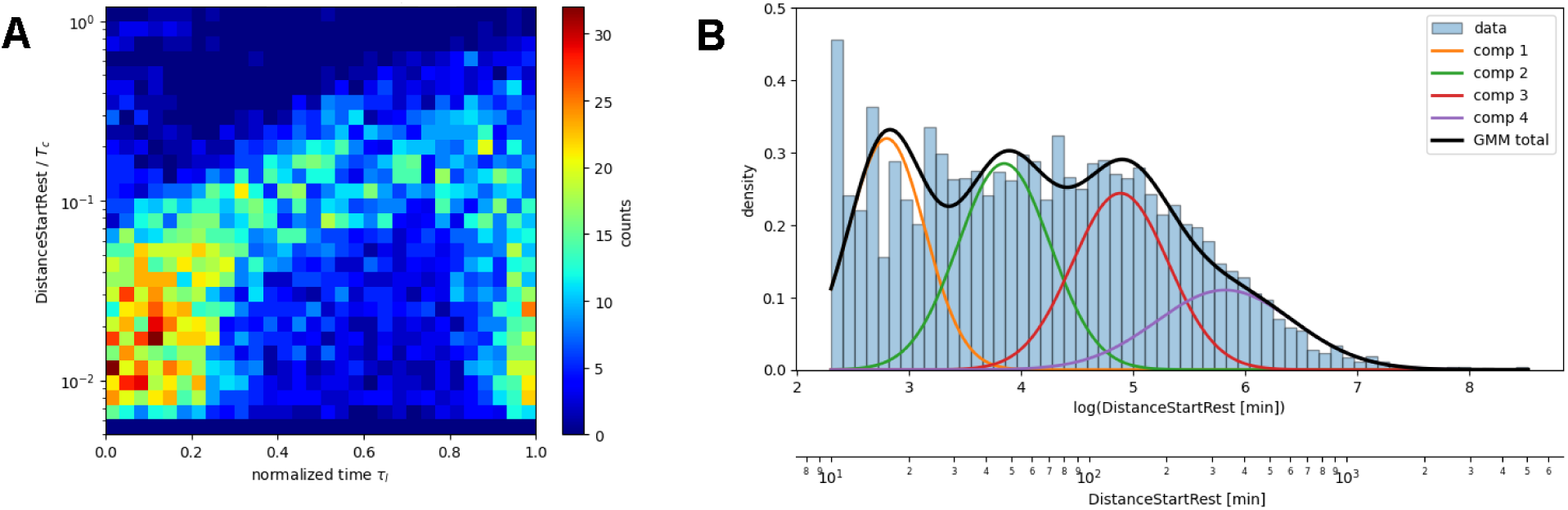
Rhythm Analysis performed with *DistanceStartRest*, high activity group. (A) Heatmap visualizing the activity by *DistanceStartRest/T_c_* against the normalized time *τ_i_*, analogous to Fig 10. (B) Gaussian Mixture Model fitted to log-transformed *DistanceStartRest* values, analogous to Fig 14.

#### 5.2.3 Temporal patterns and Gaussian Mixture Model

The clusters found in the heatmap representation of the scaled times and distances can be formalized using a K-Means cluster analysis with the variables (t, *d)* While there are several clusters with T-coordinate of the centroids less than 0.2 or larger than 0.8, their d-coordinate is always around 0.05 corresponding to *DistanceMaxActivity* ≈ 1 *hour.* There is but one in the high activity group at t ≈ 0.5 with d-coordinate *d* ≈ 0.2 corresponding to *DistanceMaxActivity* ≈ 5 *hours.* In the low activity group, this cluster is shifted to higher t values and higher *d* values. The results are in agreement with the qualitative arguments given in Section 4.1.

The parameters of the Gaussian Mixture Model are estimates based on a finite sample and have consequently an inherent statistical uncertainty. Bootstrapping can be used to quantify this uncertainty. This is a resampling method in which one approximates the sampling distribution of a statistic by repeatedly drawing samples with replacement from the empirical distribution of the observed data and recalculating the statistic on each resample [29]. Applying this method to the expectation values of the components of the Gaussian mixture model yields the error bounds given in Table 2. It can be concluded that the expectation values of the first two components of the high activity group are within the error bounds of the low activity group, whereas the expectation values of the third and the fourth components are significantly different. This again emphasizes that rest times in the low activity group are significantly longer.

**Table 2.** Errors of the components of the Gaussian Mixture Model obtained by bootstrapping.

|  | High activity group |  | Low activity group |  |
| --- | --- | --- | --- | --- |
| component | $E[X], [h]$ | Error, [h] | $E[X], [h]$ | error, [h] |
| 1 | 0.607 | 0.0873 | 0.624 | 0.012 |
| 2 | 1.41 | 0.205 | 1.52 | 0.396 |
| 3 | 3.17 | 0.0238 | 4.12 | 0.0783 |
| 4 | 7.74 | 0.456 | 14.2 | 0.67 |

### 5.3 Relevant information obtained about cockroach behavior

In this paper, behavioral data of Madeira cockroaches were interpreted as time series. It was found that the time distance between activity impulses contains valuable information on the activity behavior. Therefore, these values are ideal candidates for determining ultradian rhythms. However, there is no obvious periodic structure which can be inferred from these data in a straightforward way, but any analysis has to be based on unsupervised machine learning methods leaning strongly on stochastic models. Consequently, it is suggested that ultradian rhythms are generated by a superposition of several oscillators which are stochastic in their nature.

All analyses described in this paper detected an ultradian rhythm of the maxima of activity impulses of about 1 hour. This ultradian rhythm appears to be gated via the circadian clock since it was mostly detectable at the subjective night, the activity phase of the nocturnal cockroach. Apparently, this 1h rhythm could be split up in two components, one at about 40 minutes (0.66 hours) and the other one at about 1.5 hours. The time distance between activity impulses (metric *MaxActivity, see* Fig 3) is measured here as the time distance between maxima of activity impulses (metric *DistanceMaxActivity,* see Fig 3). This time distance can be much longer in the subjective day and may last from several hours to a complete circadian period. Activity pauses during subjective day seem to distinguish groups of animals of high activity from those of low activity. High activity animals have an average return period of the metric *MaxActivity* of about 5h hours during subjective daytime, whereas low activity animals may have a much longer return period during subjective day or show no activity at all in this time span. Fig 17 illustrates these findings with the parameters for both groups taken from the heatmaps in Fig 12.

**Fig 17.**
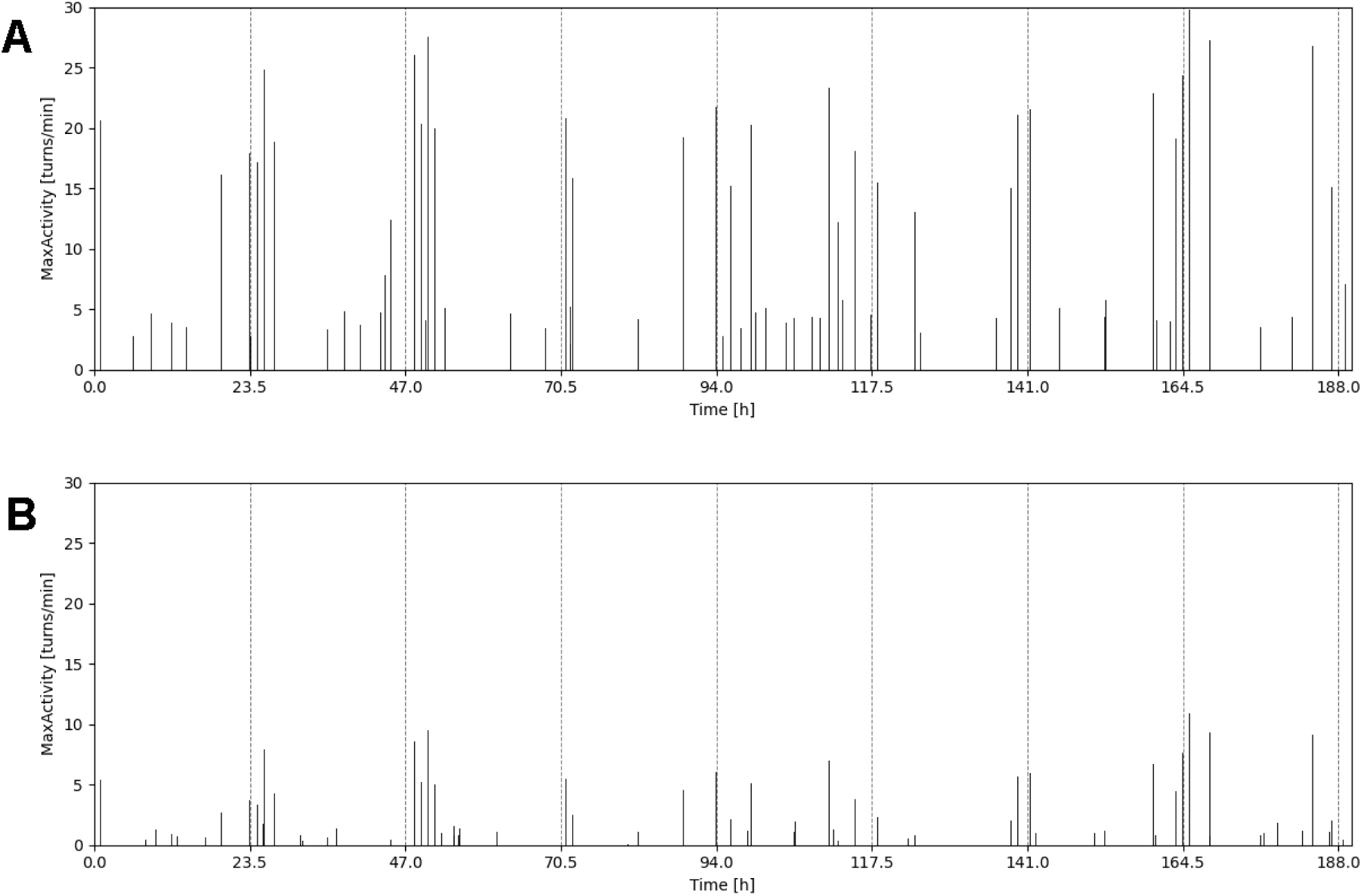
Visualization of circadian and ultradian rhythms for both activity groups. (A) High activity group. (B) Low activity group. The assumed circadian rhythm has a return period of 23.5h (dashed vertical lines), ultradian rhythms and their temporal pattern are taken from Fig 12.

The biological interpretation of the metric used in this study (the time interval between successive activity- cluster maxima) requires careful consideration. In some cases, this interval may indeed represent a recovery period following a locomotor bout. However, this interpretation is not universally applicable. During prolonged periods of activity, locomotion often remains elevated between successive maxima, with little or no quiescent interval. In such cases, the distance between maxima cannot be regarded as a simple recovery interval. Instead, the observed temporal pattern may reflect transitions between internal behavioral states rather than alternating phases of exhaustion and recovery. Depending on age, hormones, and different internal physiological states cockroaches likely switch dynamically between behavioral motor programs such as spontaneous locomotion, exploratory behavior, foraging, grooming, and rest [30–32]. Thus, activity clusters of running wheel behavior may represent periods of increased behavioral activation, while the timing of successive maxima possibly reflects the temporal organization of these state transitions.

The ultradian locomotor activity rhythm may therefore modulate the probability of initiating or maintaining locomotor activity rather than imposing discrete bouts separated by fixed recovery periods. During long-lasting activity clusters, multiple ultradian cycles may become merged into a single extended episode of locomotion, producing local fluctuations in activity without complete behavioral disengagement. Consequently, the timing of cluster maxima should not be interpreted as a direct readout of the underlying oscillator.

Our analysis therefore most likely captures the behavioral output of the endogenous ultradian oscillator rather than the oscillator itself. The rhythm identified in the sequence of activity-cluster maxima may thus serve as a marker of the temporal organization of behavior, emerging from the interaction between multiscale endogenous ultradian oscillators/clocks, variable bout durations, and stochastic behavioral decisions. Accordingly, the intervals between successive activity-cluster maxima should not necessarily be interpreted as recovery intervals but rather as reflecting the dynamics of behavioral state transitions shaped by both rhythmic and non- rhythmic influences.

## 6. Conclusions

Locomotor activity data of nocturnal Madeira cockroaches under constant conditions are analyzed in search for multiscale endogenous rhythms using methods taken from unsupervised machine learning. The following approach is taken:

- Metrics of the activity sequences are defined. The metrics are based on activity impulses, i.e. time segments of continued activity.
- Identification of distinct groups of different activity level is based on a K-Means cluster analysis.
- Characteristic features of the activity metrics are extracted using a Gaussian mixture model.

With this approach, it is possible to analyze locomotor data on a sub-circadian level, and to identify potential candidates for ultradian rhythms. A detailed study of the temporal pattern of activity reveals that the distance between activity impulses is about one hour in periods of high activity (‘night’), whereas periods of low activity (‘day’) are characterized by much lower repetition times of activity impulses ranging from about 5 hours for highly active animals to a complete circadian cycle for low activity animals. These findings suggest that there are ultradian rhythms in locomotor activity which are also linked to the circadian rhythm. However, it has to be kept in mind that these ultradian rhythms are stochastic in their nature and cannot be identified as directly as the circadian rhythm in running wheel behavior of animals. Nevertheless, it is shown that the results obtained by unsupervised machine learning are stable in the sense that changing the model assumptions does not greatly influence the final results.

